# Carbonic anhydrase network of genes trigger cytosolic pH enabling differentiation from quiescence

**DOI:** 10.1101/424648

**Authors:** Jeshima Khan Yasin, Sakshi Chaudhary, Nidhi Verma, Sengottaiyan Vennila

## Abstract

**Background:** Carbonic anhydrase regulates various cellular processes. Intracellular pH flux impacted by carbonic anhydrase alters the enzyme’s allosteric active site which effects several downstream cellular processes. Earlier, we reported that, the catalytic activity of carbonic anhydrase is independent, but direction of catalysis is affected by cellular pH level. On the other hand carbonic anhydrase alters the cytosolic pH level to facilitate allosteric phosphorylation of proteins which further leads to cellular differentiation through a process being regulated by ncRNAs.

**Results:** This study illustrates various ways of cell differentiation/ organ development regulation *via* carbonic anhydrase interacting network of proteins involved in various cellular processes. It is involved in protein degradation process of other proteins like RPT, 26S proteasome, AT3G15120 and its variant producing ncRNA, etc. Carbonic anhydrase indirectly involved in signaling process along with MAPK in providing innate resistance against biotic and abiotic stresses. It is also indirectly linked to cell membrane transporters like H^+^-ATPase and V-ATPase B.

**Conclusions:** Though carbonic anhydrase is not directly linked with EMS1 as revealed by network analyses and protein-protein interaction there could be a suitable condition generated by the carbonic anhydrase for EMS1 to be active. Hence, we report that carbonic anhydrase, along with other pH regulating gene complexes plays a major role for making EMS1 functional.

## Background

Carbonic anhydrase (CA) network of genes play a regulatory role in maintaining the intracellular cytosolic pH. Plant cell differentiation is a result of maintained pH flux influence, but detailed investigation of its impact in cell differentiation mechanism is warranted. The regulation of intracellular pH is a fundamental physiological process of great importance in growth and metabolism of every cell. Allosteric modulations of amino acids that take part in cellular differentiation are mainly altered by pH variations ranging between 6.5 and 7.5 (Minocha, 1987). There is homeostatic mechanism of pH flux within cytoplasm and cell devotes lots of effort to regulate its pH by transport of ions, hormones and nutrients for survival at the expense of energy. Thus, it has great significance in metabolism and plant growth. Non-coding class of RNAs have a crucial role in controlling gene expression in development as well as differentiation and long non-coding RNAs initiate co-expression based competitive feedback regulation of differentiation (Fatica and Bozzoni, 2014). Growth, development and cellular differentiation arise from a network of twining mechanisms. Intracellular stress is expressed and estimated in terms of oxidative stress and reactive oxygen species. Oxidative stress (OS) instigates cellular senescence, DNA damage and deleterious gene or suppressor induced loss through a cascade of gene action. OS can control miRNA interceded gene silencing in senescence, by affecting miRs expression ensuing in abounding ROS.

Thermodynamic equilibrium between CO_2_ and HCO_3_^−^ impacts the inverse relationship between rates of CO_2_ hydration and HCO_3_^−^ dehydration by carbonic anhydrase to pH dependencies, but independent of the catalytic mechanism (Yasin et al., 2016 and Yasin, 2015). CA interconversion catalysing metalloenzyme of HCO_3_^−^ and CO_2_ is a major protein component of higher plant tissues. CA exists among all kingdoms of living organisms in ☐, β, γ, δ, ε, and ζ forms without sequence homologies catalysing the reaction indicating multiple modes of gene regulation. CA converts CO_2_ to HCO_3_^−^ for phosphoenolpyruvate carboxylase reaction and convert HCO_3_^−^ to CO_2_ for ribulose-1,5-bisphosphate carboxylase reaction in photosynthesis. Moreover, CA activity in guard cells is required for CO_2_ mediated stomatal regulation and indirectly protect against stress conditions (Wei-Hong et al., 2014). Meldrum and Roughton (1934) observed that the catalytic effect of the enzyme upon the hydration was smaller at pH 10 than at pH 7.6 which indicates the pH dependence of intracellular activities.

Zinc complex facilitates carbon dioxide hydration. At pH 8, this reaction reaches a maximal rate as pH is inversely proportionate to the rate of reaction. The median is near pH 7, indicating p*K*_a_ 7 plays an important role in CA activity and the deprotonated (high pH) form of this group effects catalysis. This transition is mediated by zinc-bound water molecule. Thus, binding of water molecules to the positively charged zinc centre reduces the p*K*_a_ from 15.7 to 7. Lowered p*K*_a_ generates a substantial concentration of hydroxide ion (bound to zinc) at neutral pH. A zinc-bound hydroxide ion is nucleophilic and catalyses carbon dioxide hydration much more readily than water does (Berg et al., 2002). Lowering pH is correlated to down regulation of *cah1* transcript (Fett and Coleman, 1994). The increased CA activity at high pH appears to be required for the maintenance of the HCO_3_/CO_2_ equilibrium which is a function of the rate of CO_2_ supply available for transport. PTM (post translational modifications) of CA was proposed to activate EMS1 (Huang et al., 2017). But, we earlier reported that, CA plays major role in maintaining the intracellular pH as a key role player of pH dependencies of the cell for its intracellular functions and downstream activation of pathways (Yasin and Chaudhary, 2016; Yasin and Magadum, 2016a; Yasin and Magadum, 2016b). In this present investigation we investigated and presented the network analyses of CA with EMS1.

## Materials and methods

### Network prediction

The raw data of 19 loci with 49 gene models with a total of 64 variants were retrieved from databases including tair url (*https://www.arabidopsis.org/servlets/Search?type=general&search_action=detail&method=1&show_obsolete=F&name=carbonic+anhydrase&sub_type=gene&SEARCH_EXACT=4&SEARCH_CONTAINS=1*) and ncbi (*https://www.ncbi.nlm.nih.gov/gene*). Using these initial data sets, evidence based new network has been predicted and generated using previously reported gene function, co-expression and co-occurrence retrieved from String database (Szklarczyk et al. 2017) using String V.10 with a high confidence level of 0.7 (Szklarczyk et al. 2015). Each primary node was taken with a maximum of 10 interactions and secondary nodes with a maximum of five interactions. Disconnected nodes were removed to identify a network of 597 genes for further analyses.

### Homology modelling

BCA1 and EMS1 structure was predicted based on homology modelling web server SWISS-MODEL. SWISS-MODEL (*https://swissmodel.expasy.org/)* is a mechanised system for modelling the 3D structure of a protein (Schwede et al., 2003; Guex et al., 2009; Bordoli et al., 2009; Kiefer et al., 2009). It is user-friendly web interface to generate 3D models for their protein (Schwede et al., 2003; Arnold et al., 2006).

### Phosphorylation site prediction

Phosphorylation sites of βCA1 and EMS1 were predicted using NetPhos3.1. The NetPhos 3.1 server (*http://www.cbs.dtu.dk/services/NetPhos/*) predicts threonine tyrosine and serine phosphorylation sites in eukaryotic proteins using ensembles of neural networks. Both generic and kinase specific predictions are performed (Blom et al., 1999).

### Docking

Protein–protein interactions mediate most cellular functions; thus, a detailed description of the association process at molecular level is essential to comprehend the fundamental processes that sustain life (Ritchie, 2008). pyDOCKWEB server (http://life.bsc.es/servlet/pydock) (Jiménez-García et al., 2013) was used in the present investigation for docking of BCA1 and EMS1.

### Electrostatic properties of protein

Electrostatic effects arising due to the interaction of solute charges among themselves and with solvent and ion charges, are of utmost importance for bimolecular structure and function (Lu et al., 2008) We have predicted it using Bluues (*http://protein.bio.unipd.it/bluues/*) (Fogolari et al., 2012).

## Results

From sequence analyses 19 CA genes from arabidopsis (table 1) were identified and listed to interact with different groups of genes. Co-expression based data analyses in developing a network indicated the presence of 597 genes linked either directly or indirectly with 19 carbonic anhydrase genes in arabidopsis. The gene name, node identifier and its annotations were listed in the table. Majority of the genes listed are of energy related and transporter genes involved in intracellular and intercellular transport. PTM (post translational modifications) of β-CA is actually favoured by CA and its network (Fig. 1). Structure of EMS1 (Supplementary material 1) and β-CA were (Supplementary material 1) developed through homology modelling. Phosphorylation details of EMS1 (Fig. 2) and β-CA (Fig. 3) were illustrated in the images.

**Fig. 1.**
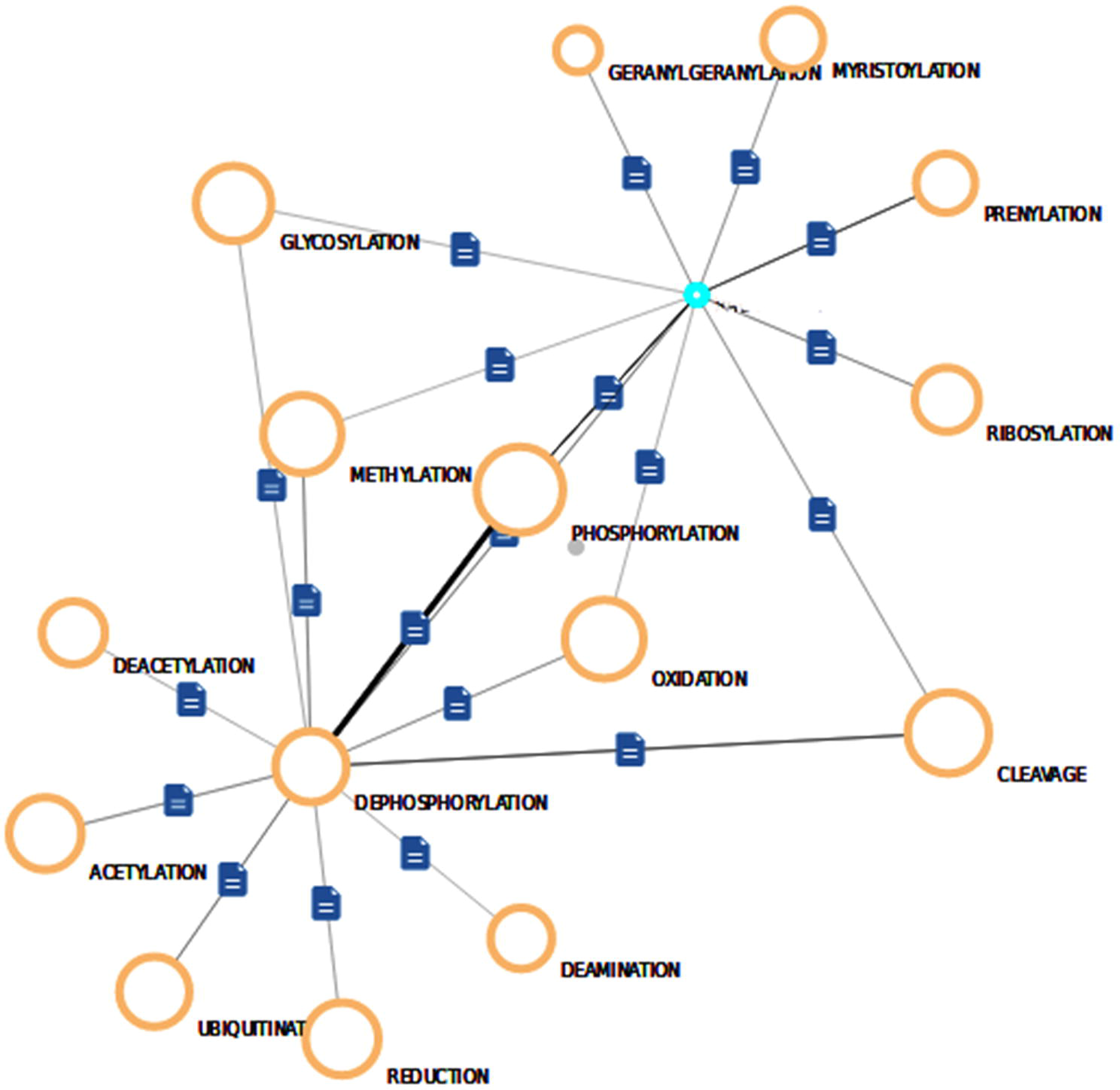
Network of post translational modifications indicating pathway of carbonic anhydrase gene cluster of *Arabidopsis thaliana*

**Fig. 2.**
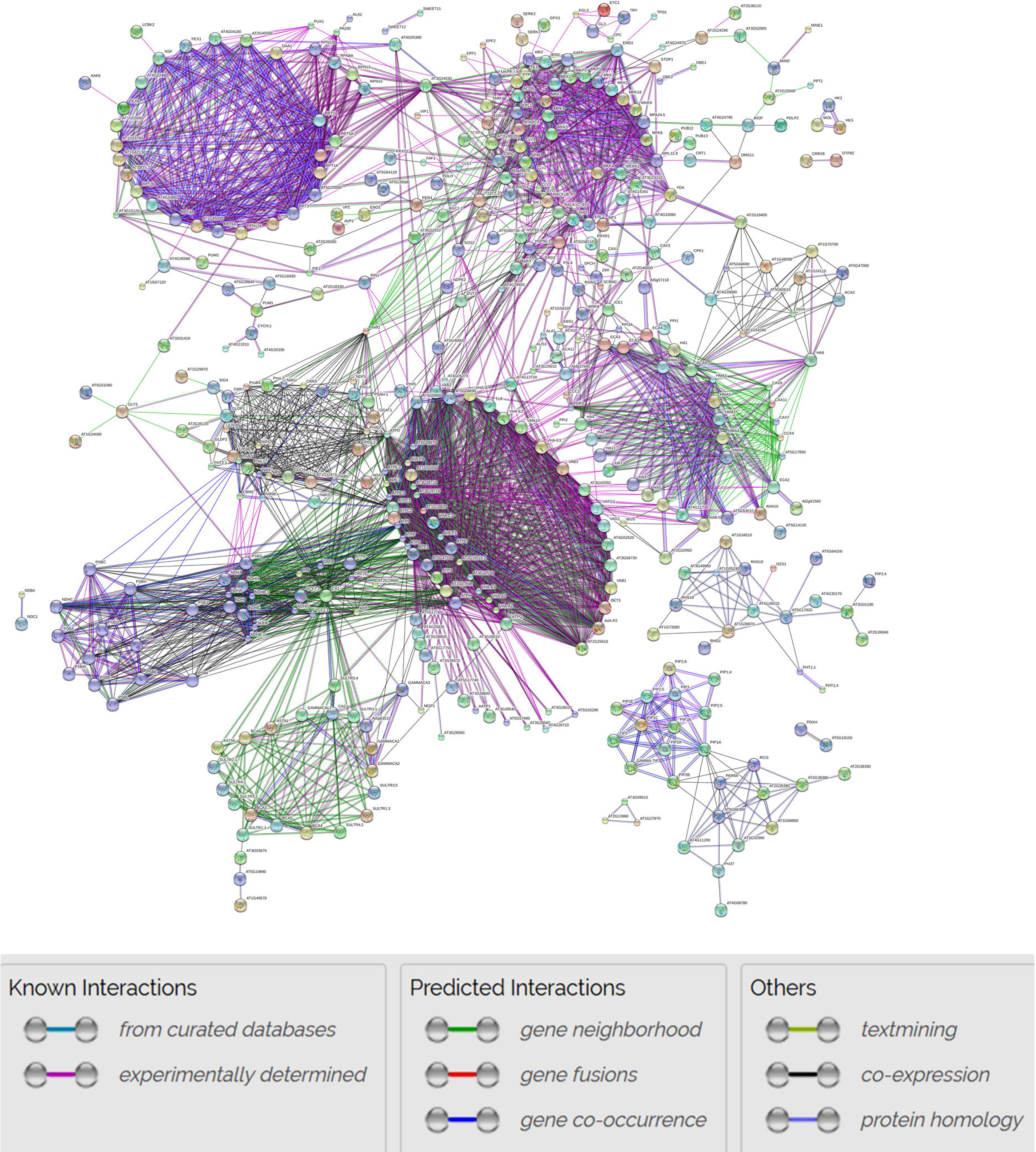
Phosphorylation sites of EMS1

**Fig. 3.**
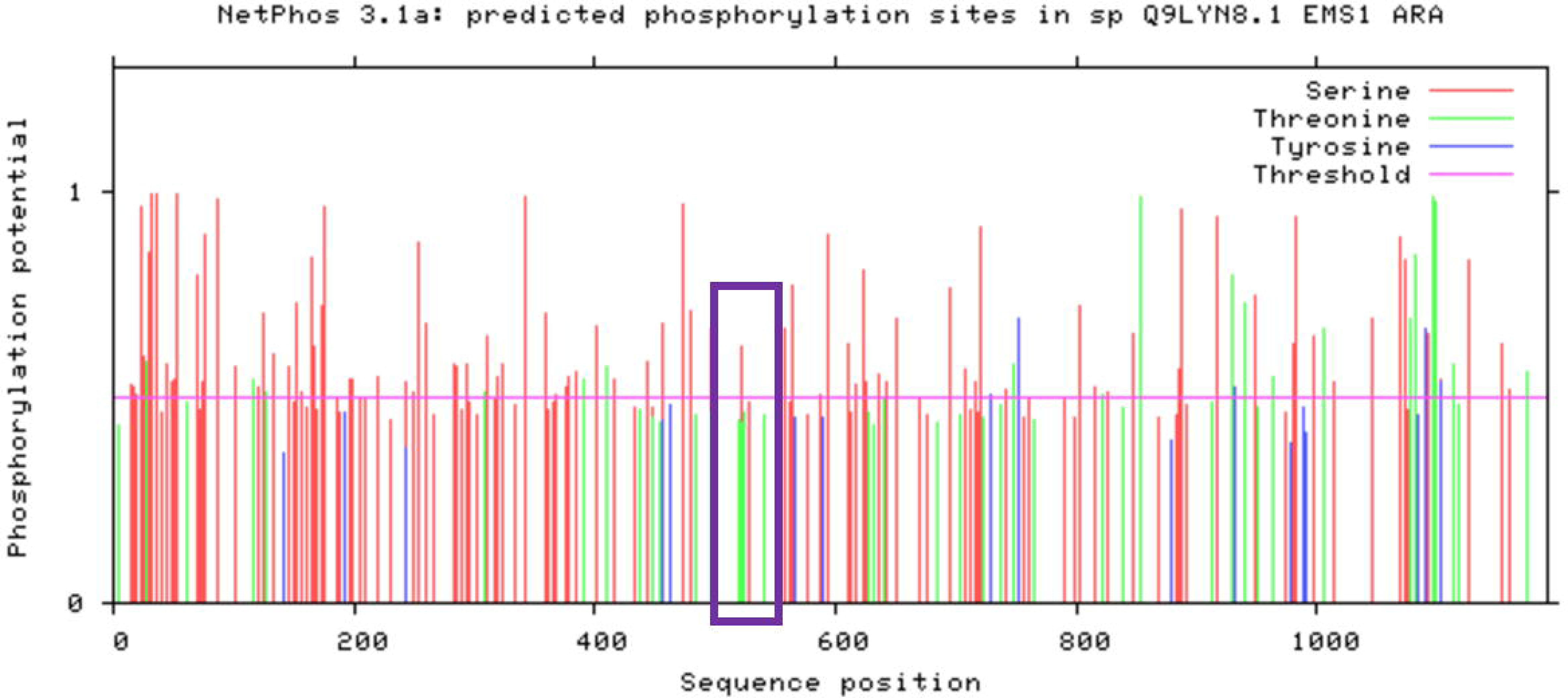
Phosphorylation sites of CA

A group of 18 ATP genes are functionally linked to CA1. Of the total 597 genes (Fig. 4 and Supplementary material 3, 4), 266 genes are energy related genes. This confirms our earlier reports of energy regulated, pH flux impacted structural compaction, as stress tolerance mechanism (Yasin et al., 2012; Yasin et al., 2014). HA cluster genes are indirectly linked to CA. HMA (Heavy metal ATPase) family plays an important role in transition metal transport in plants. It consists of five members, namely HMA1- HMA5. Total eight members of CAX (Cation exchanger /proton exchanger) are present in this network namely CAX1-7 and CAX9. This cluster is indirectly linked to CA but not with EMS1.

**Fig. 4.**
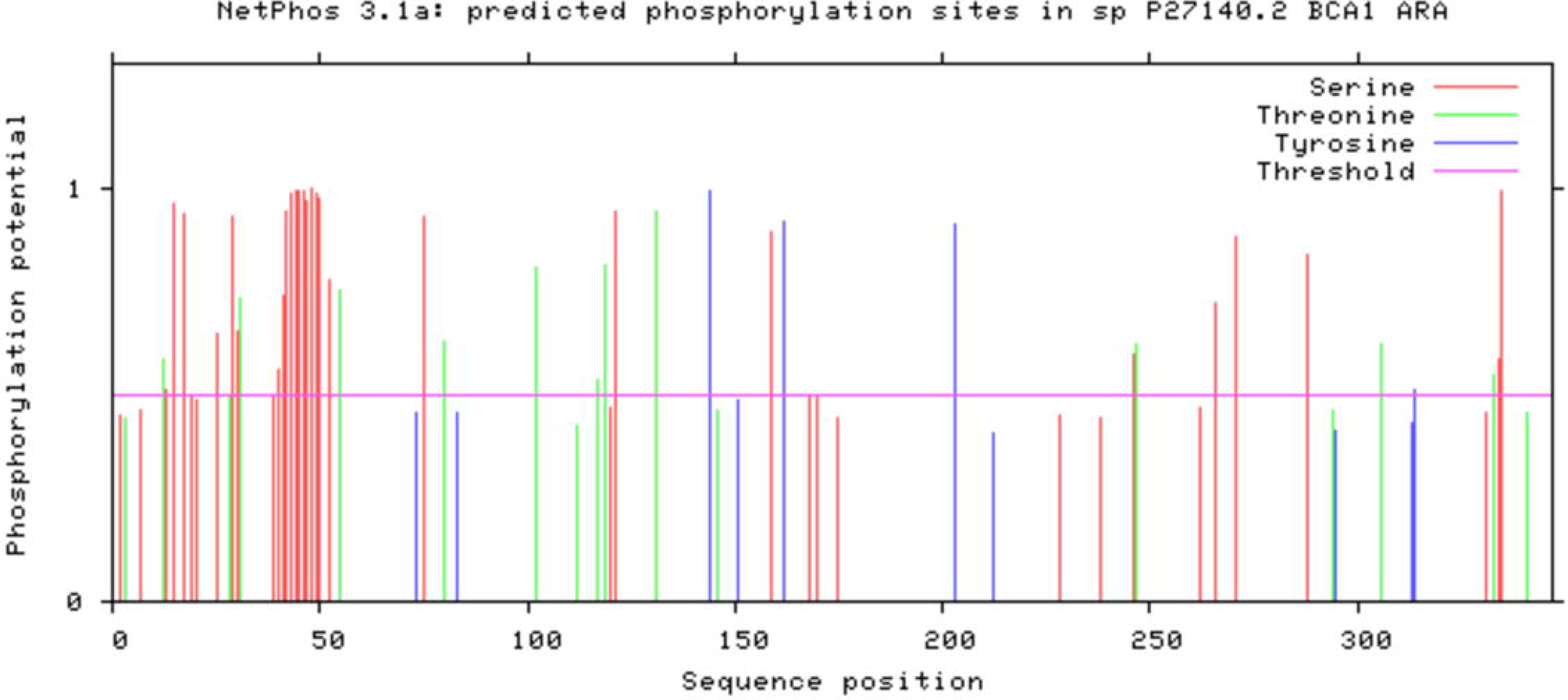
Network of genes interacting directly and indirectly with carbonic anhydrase gene cluster of *Arabidopsis thaliana*

MAPK (MAP kinase kinase) cluster of proteins consists of MKK5, MKK7, MKK3, MKK2, MKK9, MKK4, MPK3, MPK18, MPK6, MEK1, MKP2, MKP1, MPA24.5, MPL12.8, MEKK1, YDA, EMS1 and STOP1. YDA and STOP1 are the major nodes of interaction linking signalling pathway genes, innate immunity to pathogens and response to biotic and abiotic stresses. EMS1 is functionally related to ICE1, STOP1 and RIN1 indirectly. Further, these genes are indirectly connected with CA. More than 350 direct interactions of CA with other genes were identified from this network (Supplemental Material 3,4). Interaction analyses *in-silico* of CA and EMS1 revealed the absence of molecular interaction (Fig. 5,6,7 and Supplemental Material 5,6,7).

**Fig. 5.**
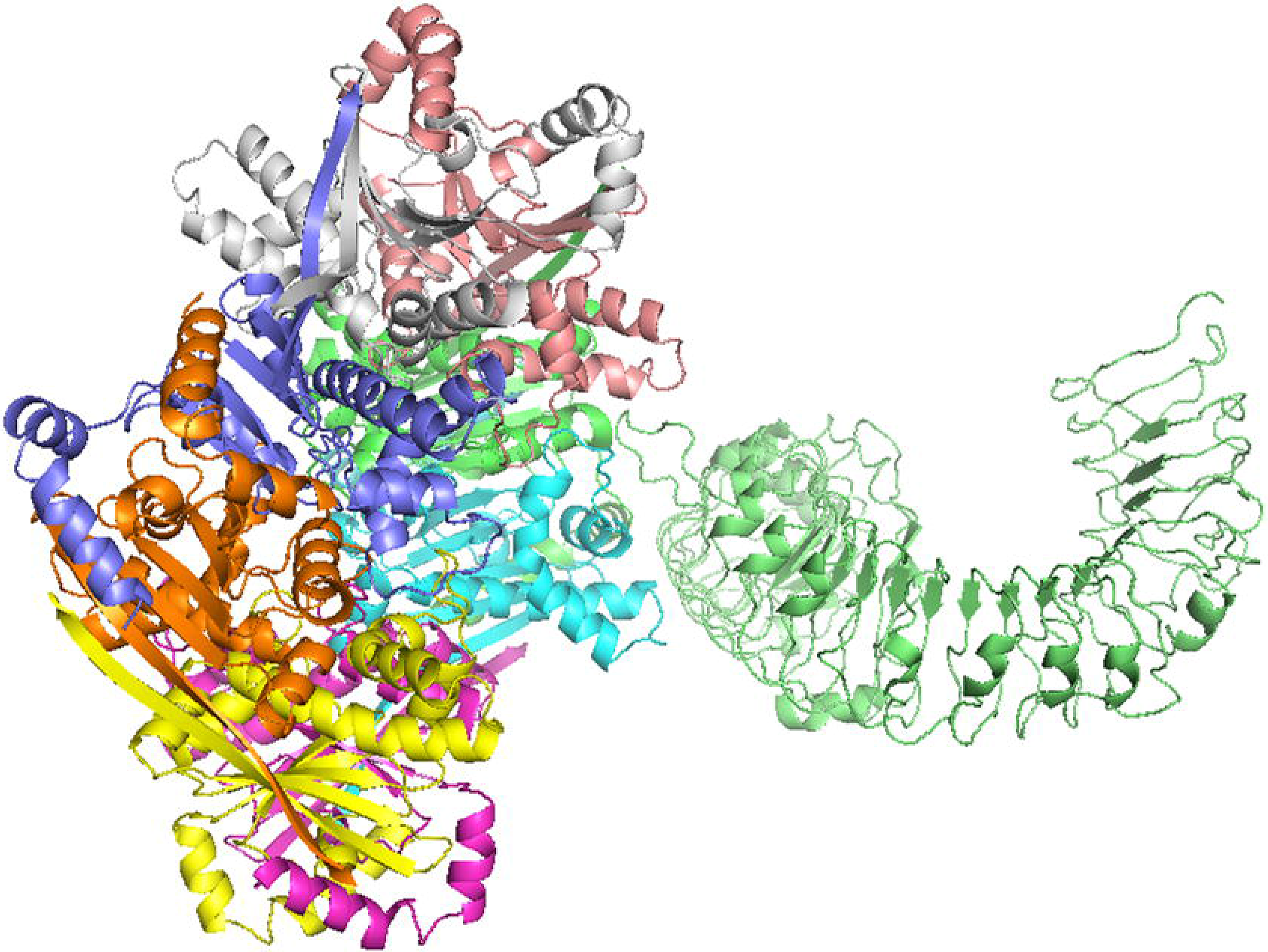
No interaction between CA and EMS1

**Fig. 6.**
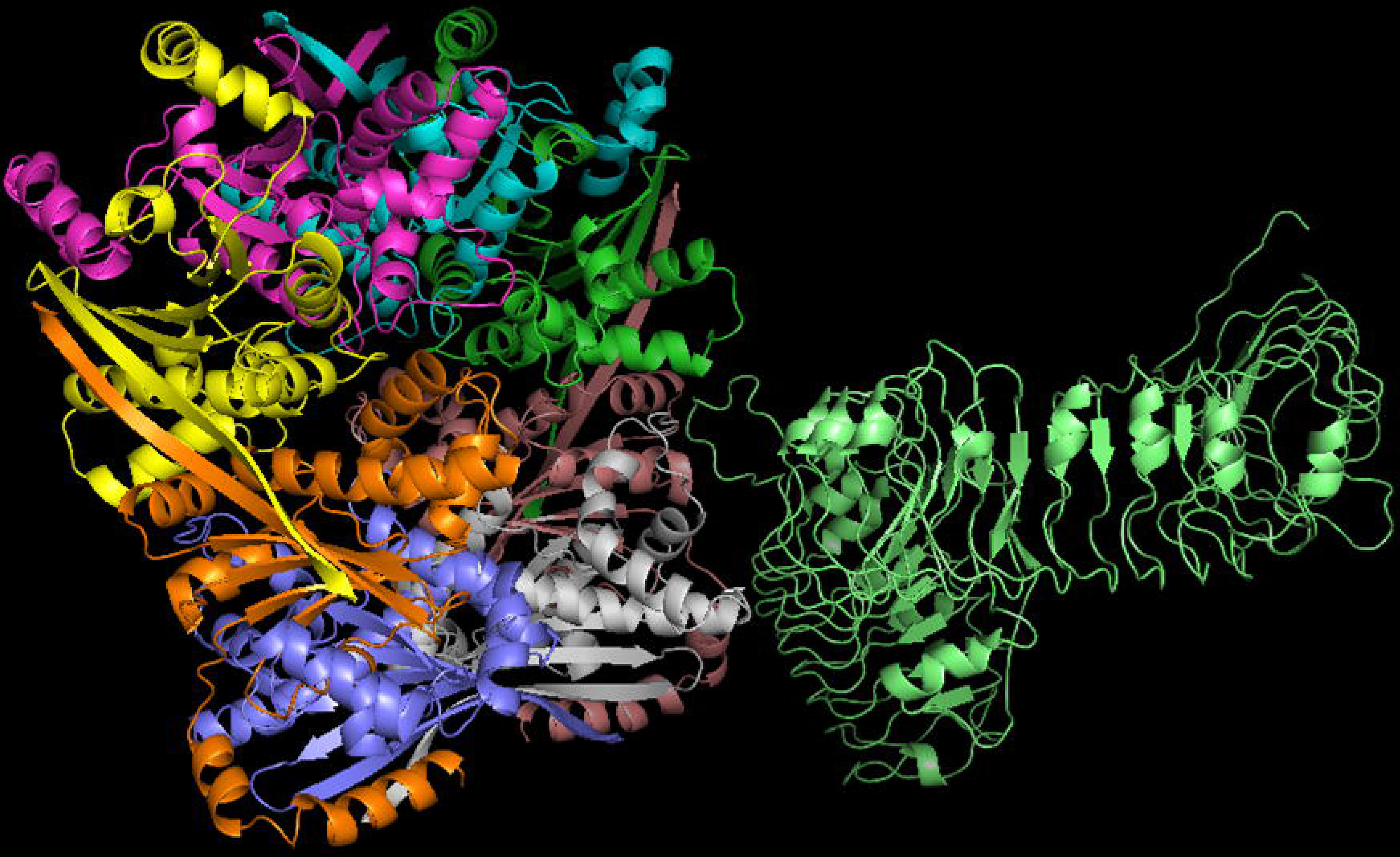
No interaction between CA and EMS1

**Fig. 7.**
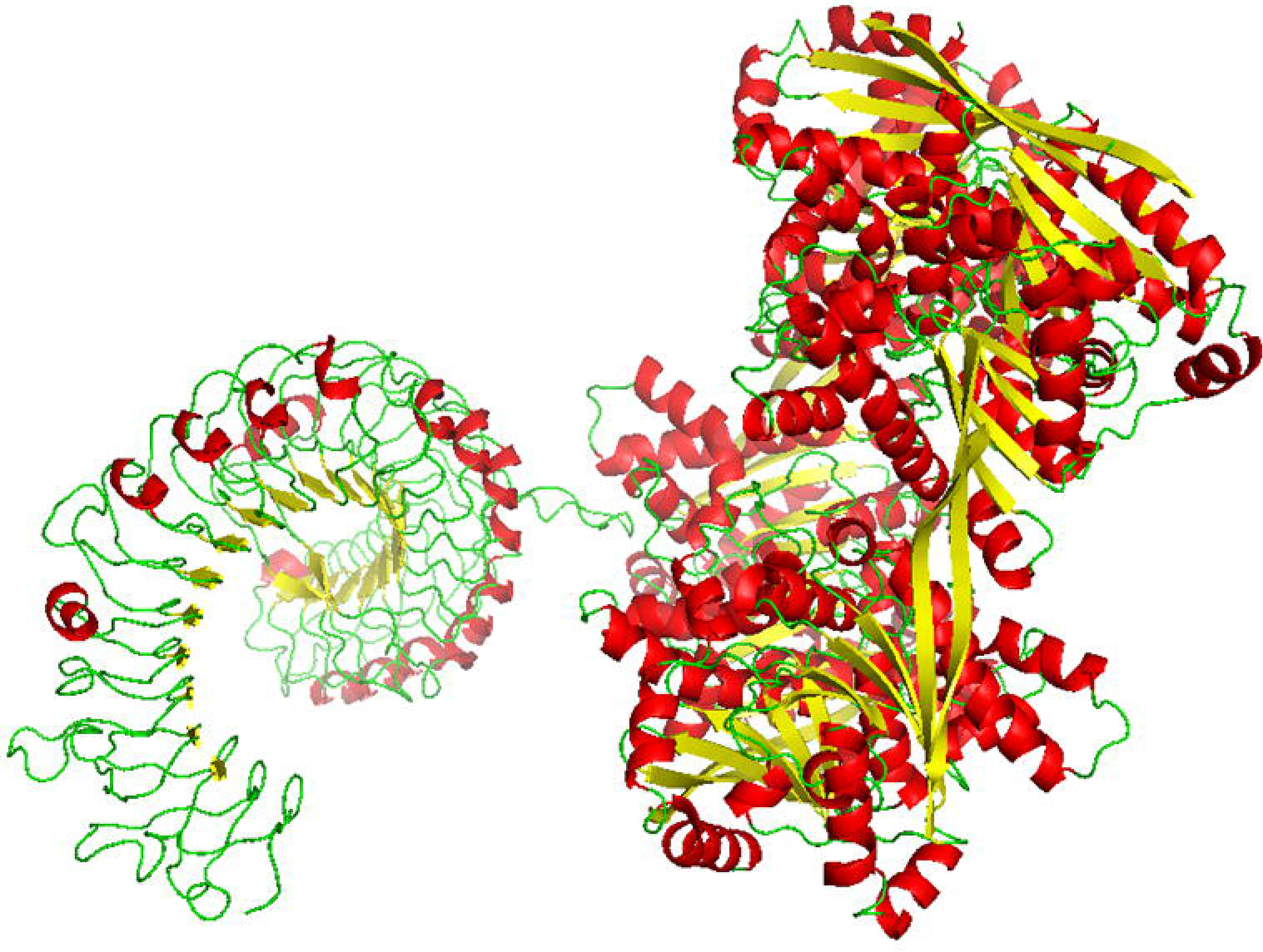
No interaction between CA and EMS1

Network of genes, indicating the co-expression interactions between proteins and small molecules essentially elucidate understanding of molecular and biochemical linkages between CA, its regulators and its impactors are presented in Supplemental Material 4 and 5. Co-expression data indicate that group of RPT (Regulatory particle triple-A) proteins consisting of RPT1A, RPT2a, RPT2b, RPT3, RPT4A, RPT5A, RPT5B and RPT6A interact with CA linked genes network. A 26S protease involved in ATP-dependent degradation of ubiquitinated protein is linked to RPTs indicating the presence of degradation pathway of gene regulation. Tight link of AT3G15120, a p-loop containing partial ncRNA splice variant generating gene confirms this mode of gene regulation. The ATPase complex confers ATP dependency and substrate specificity to the 26S complex. RPT2a (Regulatory particle AAA-ATPase 2A) is required for the maintenance of postembryonic root and shoot meristems. It has a specific role in regulation of organ size.

## Disscussion

In an earlier investigation we reported 11 carbonic anhydrase genes from *Glycine max* (Singh et al. 2017). There is a contrary link between rates of hydration and dehydration by carbonic anhydrase, which is linked to intracellular pH and transport, other than the one explained by Huang et al. (2017), which is being narrated in this manuscript. PTM is playing a major role in making any protein functional. Phosphorylation of β-CA favoured the differentiation of tapetal cells. β-CA regulates pH of tapetal cells. Huang et al. (2017) have identified a downstream pathway linking kinases with β-CA in tapetal layer differentiation. EMS1 (excess microsporocytes1) in turn initiates signaling via LRR-RLKs (leucine-rich repeat receptor-like kinases) for differentiation with co-acting SERKs, whereas, over-expression of β-CA failed to produce fertile pollen and long siliques in their experiments due to lack of pH balance required for the cell to survive and regulate through feedback inhibition. Hence, this supports our earlier findings that pH balance leads the cells to either quiescence or differentiate into a functional cell (Yasin and Chaudhary, 2016; Yasin and Magadum, 2016a; Yasin and Magadum, 2016b) which is impacted by a network of genes (Fig. 2). β-CA in steady state kinetics tangle within protein allosteric site alteration. In many instances β-CA was reported to be activated by pH alteration and resulting allosteric modulation leads altered cellular growth (Lindskog and Coleman, 1973). Whereas, CA family contributes to intracellular pH and its balance leads to allosteric changes and resulting cellular activities.

RPT5A plays an essential role in the gametophyte development, but not linked to CA directly. RPN10 plays a regulatory role for 19S regulatory particle (RP) of 26S proteasome (Fu et al., 1999) and recognises ubiquitinated substrates for ubiquitin/26S proteasome-mediated proteolysis (UPP) either directly or indirectly and selects ubiquitin-conjugates for destruction. RPN10 prefers multiubiquitin chains rather than single ubiquitins and acts as a potential docking subunit for both ubiquitin receptors RAD23s and DSK2s, with a binding affinity for ‘Lys-48’-linked ubiquitin chains. RPN12a, RPN12b acts as a regulatory subunit of the 26S proteasome in ATP-dependent degradation of ubiquitinated proteins. RPN 13 functions as a proteasomal ubiquitin receptor with a binding affinity for ‘Lys-63’-linked ubiquitin chains. PEX genes are involved in protein import into peroxisomes, recycling, jasmonate biosynthesis, developmental elimination of obsolete peroxisome matrix proteins and may form heteromeric AAA ATPase complexes required for the import of proteins.

Genes from this cluster contribute to stomatal cell fate, regulate guard mother cell (GMC), secondary hydrogen peroxide generation during hypersensitive response-like cell death, negative regulation of polar auxin transport, positive regulation of plant basal and systemic acquired resistance (SAR), activation of EIN3, leading to ethylene signalling (Ouaked et al., 2003), ethylene, salt stress (Liu et al., 2007) and camalexin biosynthesis (Xu et al., 2008). CLV genes are ligand-receptors of extracellular signals regulating shoot and root meristem maintenance.

H^+^-ATPase is a plasma membrane bound ATPase, generating a proton gradient that drives the active transport of nutrients by H^+^-symport. The resulting external acidification or pH flux may mediate growth responses. It consists of ten HA family members vice HA1-HA10 (Robert et al., 2000).

VAB (V-ATPase B), non-catalytic subunit of the peripheral V1 complex of vacuolar ATPase is responsible for acidifying a variety of intracellular compartments in eukaryotic cells (Ma et al., 2012). Three members of vacuolar ATPase proteins are present in this network. VHA (Vacuolar proton ATPase), a multimeric enzyme catalysing the translocation of protons across the membranes has six members in this network. ATPase6 is a mitochondrial membrane ATP synthase produces ATP from ADP in the presence of a proton gradient across the membrane which is generated by electron transport complexes of the respiratory chain. ATPE encodes epsilon subunits of chloroplast ATP synthase. It also produces ATP from ADP in the presence of a proton gradient across the membrane. F-type ATPases produces ATP from ADP in the presence of a proton or sodium gradient. V^−^ATPase is responsible for acidifying a variety of intracellular compartments in eukaryotic cells. DEETIOLATED 3 is the subunit of the peripheral V1 complex of vacuolar ATPase, responsible for acidifying a variety of intracellular compartments in eukaryotic cells.

CRR (Chlororespiration reduction) genes are required for formation, assembly or stabilization and activity of the chloroplast NAD(P)H dehydrogenase (NDH) complex of the photosynthetic electron transport chain. NDH dehydrogenase shuttles electrons from NADPH via FMN and iron-sulphur centers to quinones in the photosynthetic chain and possibly in a chloroplast (Takbayashi et al., 2009). PNS proteins shuttles electrons from NAD(P)H to quinones in the photosynthetic chain and possibly in a chloroplast respiratory chain. The immediate electron acceptor plastoquinone, couples the redox reaction to proton translocation, and thus conserves the redox energy in a proton gradient. Nine members are present in this network.

GAMMACA are the key role players with zinc-containing metalloenzymes. GAMMACA are enzymes involved in the catabolism of H_2_CO_3_ (Parisi et al., 2004). CA cluster is directly linked to members of energy cluster genes, indicating the presence of regulatory function. The earlier report on the CA network in arabidopsis and soybean indicated energy related mechanisms (Singh et al., 2016) which are substantiated by the present investigation indicating CA tightly linked to ATP genes. CAs mediates complex I assembly in mitochondria and respiration. SULTR, the sulphate transporter protein family is responsible for high-affinity H^+^ /sulphate co-transport that mediate the uptake of the environmental sulphate by plant roots under low- sulphur conditions. It plays a central role in the regulation of sulphate assimilation.

In contrary to the findings from Huang et al. (2017) and Li et al. (2017) CA facilitating allosteric phosphorylation of proteins indirectly leads to differentiation through altering the cytosolic pH the process of which is being regulated by ncRNAs. Symplasmic communication in a cell is interconnected with pH regulation and allosteric modification indirectly effected by lncRNAs as well as miRs hinders the process of cell differentiation (Badger, 2003). These regulatory molecules impact growth patterns by altering protein degradation, signal transduction, pathogen invasion and response to environment stress.

## Conclusions

For RPN linked gene network, rewiring is possible through alteration of AT3G24530 which is an AAA-type ATPase family protein. This protein effects metabolic processing and gene is linked with MAP kinase cascade cluster. Likewise STOP1 (sensitive to proton rhizotoxicity) a Cys2/His2 type zinc-finger type transcription factor and EMS1 (Excess Microsporocytes 1- a LRR receptor protein kinase) can be modified to evaluate its impact. But EMS1 is no way directly linked to CA in the presently studied network. As depicted in fig.1 phosphorylation activates any enzyme and dephosphorylation leads to ubiquitination pathway. AT3G15120, a p-loop containing nucleoside triphosphate hydrolases super family protein, is the outer link which connects this RPT network with other clusters. A p-loop containing sequence normally codes for phasiRNA through which ncRNA mediated gene regulation is effected. Thus, this gene marks the link for network rewiring option by altering its expression levels.

Rewiring of pnsb1 (chloroplastic Photosynthetic NDH subunit of subcomplex B1) could alter the CRR genes. This is linked to photosystem II cluster, ATP chains and cluster4; may alter the phenotype to a larger extent. Altering the expression of HAB (Hypersensitive to ABA) may manipulate the genotype to a larger extent in modifying the expression of CAX (calcium exchanger protein) genes. GLY3 (Glyoxalase II 3), YDA (mitogen activated protein kinase kinase kinase YODA), ATPD (F-type ATPase acting in the presence of a proton or sodium gradient) and RIN1 (Ripening inhibitor) are the minor genes of importance for network rewiring alternatives. Whereas we couldn’t see any tightly linked impact from SERK genes which is an outlier of the predicted network and is not liked to CA. SULTR3 (chloroplast localised sulphate transporter) and pnsl (chloroplastic photosynthetic NDH subunit of luminal location 5) gene cluster were found to be directly linked with CA. Production of intracellular energy, proton gradient and CA are closely packed, indicating that the pH of the cell is directly playing a major role in overall wellbeing of individual cells. Though CA is not directly linked with EMS1 as revealed by network analyses, there could be a suitable condition generated by the CA for EMS1 to be active. Hence, we report that CA, along with other pH regulating gene complexes plays a major role for making EMS1 functional.

## List of abbreviations

Atom: constituent atom in protein
ATPD-F-type: ATPase acting in the presence of a proton or sodium gradient
CA: Carbonic anhydrase
CAX: Cation exchanger /proton exchanger
CAX: calcium exchanger protein genes
Conf.: confirmation
CRR: Chlororespiration reduction
Desolv.: Dissolving power
Ele: Electrostatic potential
EMS1: excess microsporocytes1
GBR: Generalised bond radii
GLY3: Glyoxalase II 3
GMC: guard mother cell
HAB: Hypersensitive to ABA
HMA: Heavy metal ATPase
LRR-RLKs: leucine-rich repeat receptor-like kinases
MAPK-MAP: kinase kinase
OS: Oxidative stress
pKa: pH of molecule
*pnsl*chloroplastic: photosynthetic NDH subunit of luminal location 5 gene
PTM: Post translational modification
Res: residue; Chain identifier
Resn.: residue number
RIN1: Ripening inhibitor
ROS: Reactive oxygen species
RP: 19S regulatory particle of 26S proteasome
RPT2a: Regulatory particle AAA-ATPase 2A
RPT: Regulatory particle triple-A
SAR: systemic acquired resistance
STOP1: sensitive to proton rhizotoxicity a Cys2/His2 type zinc-finger transcription factor pnsb1-chloroplastic Photosynthetic NDH subunit of subcomplex B1
SULTR: sulphate transporter responsible for high-affinity H^+^ /sulphate co-transport
UPP: ubiquitin/26S proteasome-mediated proteolysis
VAB: V-ATPase B
VDW: Van der walls forces
VHA: Vacuolar proton ATPase
YDA: mitogen activated protein kinase kinase kinase YODA

## Declarations

### Acknowledgements

We are grateful to Dr. Kuldeep Singh, Director, ICAR-NBPGR for providing facilities to conduct this research.

### Availability of data

The datasets processed in this study are bundled and made available with the manuscript as supplementary information.

### Author contributions

YJK designed and planned the research; YJK and SC carried out *in-silico* analyses, SC and YJK developed graphical representations, SV and YJK compiled the results, wrote the manuscript, and YJK communicated.

### Additional Information

Supplementary information accompanying this paper will be available in the online version.

### Consent for publication

Not applicable.

### Ethics approval and consent to participate

Not applicable.

Supplemental Material 1- Structure of EMS1

Supplemental Material 2- Structure of CA

Supplemental Material 3- List of genes interacting either directly or indirectly with carbonic anhydrase genes of *Arabidopsis thaliana*

Supplemental Material 4- List of gene interactions with carbonic anhydrase genes of *Arabidopsis thaliana*

Supplemental Material 5- Interaction analyses of CA and EMS1

Supplemental Material 6- Molecular structure analyses of CA

Supplemental Material 7- Molecular structure analyses of EMS1

Atom – constituent atom in protein; Res – residue; Chain identifier; Resn. – residue number; pKa- pH of molecule; VDW – Van der walls forces; Ele- Electrostatic potential; Desolv. – Dissolving power; Conf. – confirmation; GBR- Generalised bond radii

